# Cooperative Peripheral CB1 Inhibition by Extracellular Antibody Binding and Orthosteric Antagonism

**DOI:** 10.64898/2026.09.26.754614

**Authors:** Yuan Gao, Jing Wang, Yali Lai, Hong Shan, Yang Yue, Yuxin Bai, Jingjing Hou, Junrui Li, Xinheng He, Xinyue Zhang, Yinshan Tang, Wen Hu, H. Eric Xu, Fei Xu, Heng Liu

## Abstract

The cannabinoid receptor CB1 is an important therapeutic target for obesity and metabolic disorders, yet centrally acting CB1 antagonists are limited by neuropsychiatric side effects. Although the anti-CB1 antibody Nimacimab has advanced to Phase 2 clinical trials for metabolic disorders, the molecular mechanisms underlying peripheral CB1 inhibition remain poorly understood. Here, we report cryo-EM structures of human CB1 bound to Nimacimab alone and in complex with the orthosteric antagonists Rimonabant and JD5037. Unlike previously characterized inhibitory GPCR antibodies that block ligand access through steric occlusion or directly engage orthosteric sites, Nimacimab functions through extracellular allosteric inhibition by stabilizing an inactive-like extracellular architecture of CB1. Antibody binding restricts extracellular conformational dynamics and allosterically stabilizes inactive-state signaling microswitches. Importantly, extracellular antibody engagement is structurally compatible with orthosteric antagonism and cooperatively enhances CB1 inhibition. Structural analysis further reveals how JD5037 remodels the extracellular vestibule to achieve peripheral-selective antagonism. Together, these findings establish extracellular allosteric inhibition as a distinct mechanism of GPCR regulation and provide a structural framework for the development of peripherally selective CB1 therapeutics.

## Main

Cannabinoid receptor 1 (CB1) is a class A G protein-coupled receptor (GPCR) that plays a central role in regulating energy balance, appetite, and metabolic homeostasis^1–4^. Pharmacological inhibition of CB1 has therefore long been pursued as a therapeutic strategy for obesity and metabolic disorders^5–7^. Among the inhibitors developed to date, Rimonabant represents the canonical example of this therapeutic strategy^8,9^. Rimonabant was approved for the treatment of obesity and overweight in patients with cardio-metabolic risk factors^10^, demonstrating significant efficacy in reducing body weight and improving metabolic outcomes^11^. However, it was withdrawn from clinical use because of severe neuropsychiatric adverse effects, including depression, anxiety, and suicidal ideation, attributed to its penetration into the central nervous system (CNS) and blockade of central CB1 signaling^5,9,10,12–16^. These setbacks have highlighted the critical need for therapeutic modalities that selectively suppress peripheral CB1 signaling while sparing central receptor function^6,17–19^.

Substantial efforts have focused on the development of peripherally restricted CB1 inhibitors with limited brain exposure. Among these, JD5037, a Rimonabant-derived antagonist currently in preclinical development, has emerged as a prototypical peripherally restricted CB1 inhibitor^20^. JD5037 exhibits minimal CNS penetration while retaining robust metabolic efficacy, including reduction of diet-induced weight gain, improvement of insulin sensitivity, and restoration of leptin responsiveness in preclinical models^19,21,22^. These findings support the therapeutic potential of orthosteric CB1 antagonism selectively confined to peripheral tissues. However, despite extensive pharmacological characterization, the structural basis by which JD5037 engages CB1 and the molecular determinants underlying its peripheral restriction remain unknown.

In parallel with small-molecules, antibody-based modulation of GPCRs has emerged as an important strategy to achieve peripheral selectivity, as antibodies inherently lack blood-brain barrier permeability^23–29^. Nimacimab is a first-in-class and currently the only clinically advanced peripheral CB1 inhibitory antibody, representing a distinct biologic strategy for selectively suppressing peripheral CB1 signaling for obesity and related metabolic indications^30,31^. By selectively suppressing peripheral CB1 signaling, Nimacimab improves metabolic outcomes while avoiding neuropsychiatric side effects^30,32,33^. However, the mechanism by which Nimacimab inhibits CB1 remains unresolved. In particular, whether Nimacimab functions through orthosteric occlusion, negative allosteric modulation, antagonism, or inverse agonism, remains unknown. More broadly, peripherally restricted small molecules and extracellular antibodies represent fundamentally distinct strategies for CB1 inhibition, yet whether these mechanisms are mutually exclusive or functionally cooperative remains unclear.

Here, we report cryo-EM structures of human CB1 in complex with Nimacimab alone and in combination with the orthosteric antagonists Rimonabant and JD5037. Together with structural analyses, molecular dynamics simulations, and functional assays, our study reveals that Nimacimab acts as an extracellular inverse agonist by engaging ECL2 and adjacent extracellular regions to stabilize a distinct inactive-like extracellular architecture of CB1. In contrast, Rimonabant and JD5037 suppress receptor signaling through canonical orthosteric inhibition within the transmembrane pocket. Importantly, these two inhibitory mechanisms are structurally compatible and exhibit cooperative suppression of CB1 signaling, with antibody engagement further enhancing orthosteric inhibition through coordinated stabilization of the extracellular vestibule and inactive receptor conformations. Our findings define the molecular epitope and inhibitory mechanism of a clinically relevant peripheral CB1 antibody, elucidate the structural basis of a prototypical peripherally restricted orthosteric antagonist, and establish a structural and mechanistic framework for extracellular allosteric inhibition of GPCR signaling and the rational development of peripherally selective CB1 therapeutics.

## Results

### Pharmacological characterization of CB1 inhibitors

CB1 primarily couples to Gi proteins and β-arrestin pathways to regulate downstream signaling, and inhibition of these pathways underlies the therapeutic effects of CB1 antagonists in metabolic disorders (Fig. 1a). Rimonabant and JD5037 are well-established small-molecule CB1 antagonists/inverse agonists whose inhibitory mechanisms have been extensively characterized^8,34^. In contrast, although the anti-CB1 monoclonal antibody Nimacimab has shown promising metabolic efficacy, the molecular and pharmacological basis by which it suppresses CB1 signaling remains poorly understood. To systematically define the pharmacological profiles of the CB1-targeting agents used in this study, we evaluated their effects on Gi-mediated signaling and β-arrestin recruitment using cAMP GloSensor and arrestin recruitment assays, respectively (Fig. 1b-h).

**Fig. 1.**
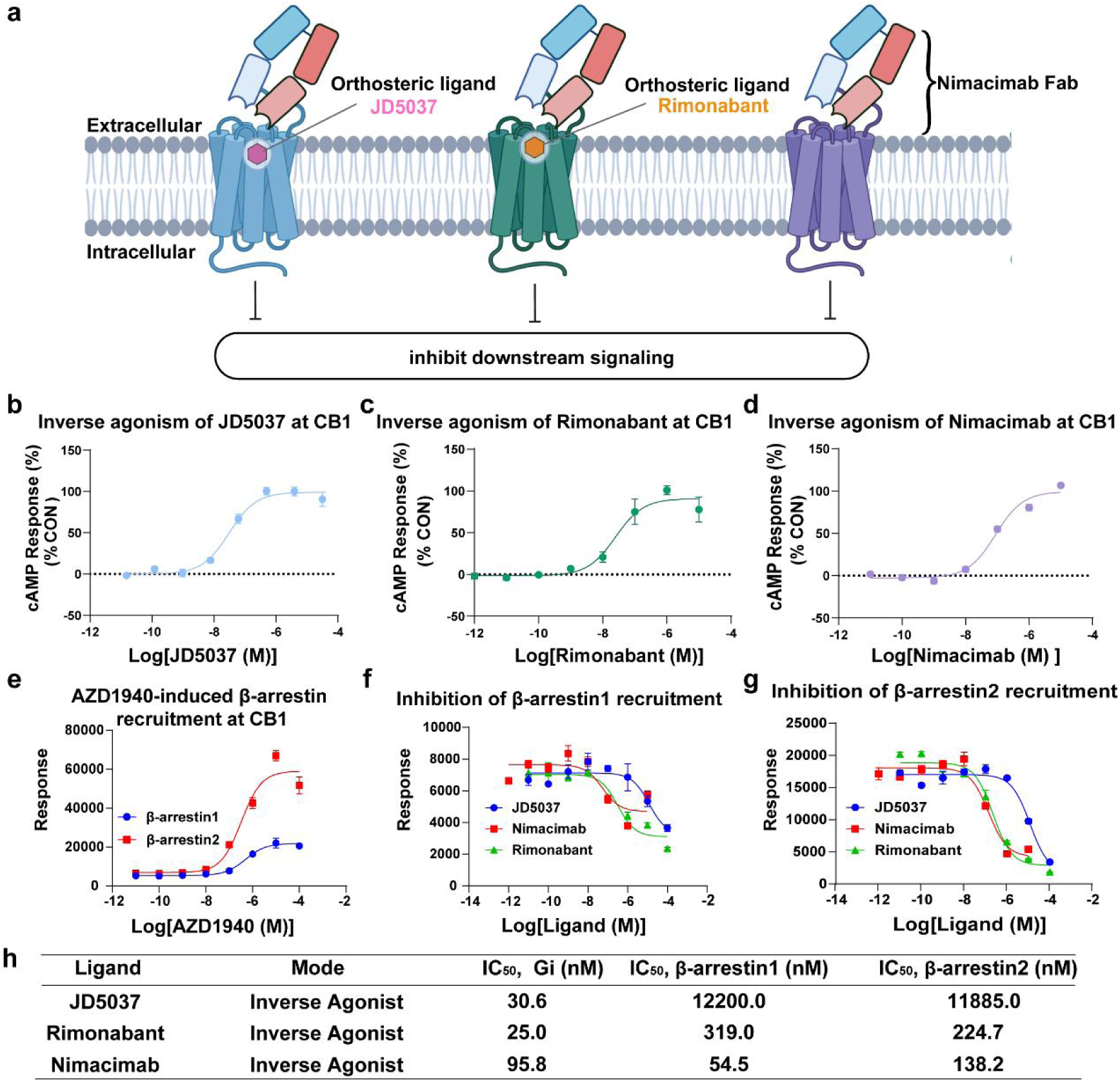
Pharmacological characterization of CB1 inhibition by Nimacimab, JD5037, and Rimonabant. a, Schematic illustration of the inhibitory modes of Nimacimab, JD5037, and Rimonabant at CB1. All three inhibitors suppress CB1-mediated downstream signaling. **b–d,** Concentration-response curves showing the inhibitory activities of JD5037**(b)**, Rimonabant**(c)**, and Nimacimab**(d)** at CB1, as measured by the cAMP GloSensor assay in CB1-expressing HEK293T cells. **e,** Concentration-response curves of AZD1940-induced β-arrestin1 and β-arrestin2 recruitment at CB1. **f and g,** Concentration-dependent inhibition of AZD1940-induced β-arrestin1**(f)** and β-arrestin2**(g)** recruitment by JD5037, Nimacimab, and Rimonabant. **(h)** Summary of inhibitory potencies of Nimacimab, JD5037, and Rimonabant in Gi-mediated cAMP signaling and β-arrestin recruitment assays. pIC_50_ values are provided in Supplementary Table 2. All functional data are shown as mean ± s.e.m. from three independent experiments (n = 3). Each independent experiment included technical replicates.

We found that both JD5037 and Rimonabant suppressed basal CB1 activity in the absence of agonist stimulation, indicating inverse agonist properties under the assay conditions (Fig. 1b, 1c, 1h and Supplementary Table 2). Notably, Nimacimab also markedly decreased basal Gi signaling, exhibiting clear inverse agonist behavior with a pIC_50_ of 7.02 (Fig. 1d, 1h and Supplementary Table 2). This inhibitory effect was achieved despite the large molecular size of the antibody and its presumed extracellular mode of receptor engagement, suggesting that CB1 activity can be suppressed independently of direct orthosteric pocket occupancy.

As for β-arrestin recruitment, our results confirmed that AZD1940 robustly induced β-arrestin1 and β-arrestin2 recruitment (Fig. 1e). We then tested inhibitor effects of ligands on β-arrestin recruitment. Rimonabant inhibited both β-arrestin1 and β-arrestin2 recruitment, although its potency toward arrestin pathways was approximately one order of magnitude lower than that observed for Gi inhibition (Fig. 1f and 1h). By comparison, Nimacimab potently suppressed both β-arrestin1 and β-arrestin2 recruitment while simultaneously inhibiting Gi signaling, demonstrating broad inhibition across major downstream CB1 pathways (Fig. 1g-h). Together, these findings establish Nimacimab as an inverse agonist capable of suppressing CB1 signaling through multiple downstream effectors, thereby providing a pharmacological framework for subsequent structural analyses.

### Overall structures of Nimacimab–CB1 complexes

To investigate the structural mechanisms of Nimacimab and orthosteric antagonism, we resolved cryo-EM structures of Nimacimab–CB1 alone and in ternary complexes with JD5037 or Rimonabant (Fig. 2a-d). To ensure high protein stability and sample quality, we employed an engineered human CB1 construct featuring a BRIL insertion in ICL3 and a series of thermostabilizing mutations as previously reported and indicated in *Methods*. This enabled the determination of four structures at resolutions of 2.7 Å–3.2 Å, the high-quality density maps permitted unambiguous atomic model building and detailed analysis of the intermolecular interactions (Extended Fig. 1-4).

**Fig. 2.**
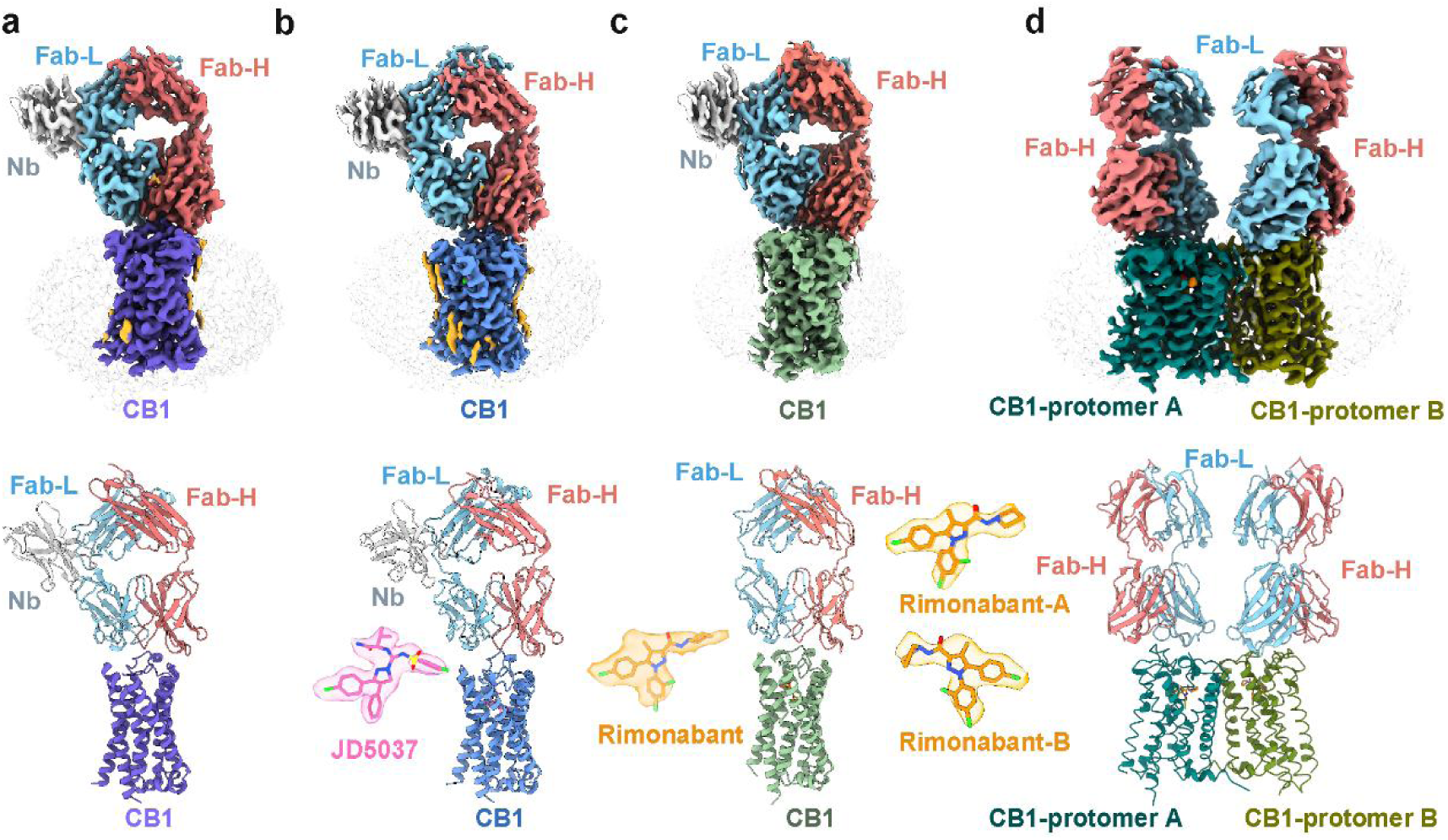
Overall cryo-EM structures of Nimacimab-bound CB1 complexes. a–d, Cryo-EM density maps and atomic models of the Nimacimab–CB1**(a),** Nimacimab–JD5037–CB1**(b),** one protomer of the Nimacimab–Rimonabant–CB1 dimer**(c)** and the full dimeric Nimacimab–Rimonabant–CB1 complex**(d).** Fab heavy chain, salmon; Fab light chain, cyan; nanobody, gray; CB1 molecules, distinct colors; JD5037, pink; Rimonabant, orange.

In all structures, the Fab fragment of Nimacimab binds to the extracellular surface of CB1 above the transmembrane domain, with the complementarity-determining regions (CDRs) primarily engaging the extracellular loops (Fig. 2a-d). The overall binding orientation of the antibody is highly conserved across all complexes, indicating a stable and ligand-independent mode of extracellular recognition. Notably, antibody engagement does not induce detectable changes in the binding mode of either small molecule, and no steric conflicts are observed between the Fab fragment and the orthosteric ligands. Instead, in the ternary complexes, the N-terminal region of CB1 adopts a lid-like conformation that encloses the orthosteric pocket and shields the bound antagonists from the extracellular environment (Extended Fig. 5a-c). These observations demonstrate that extracellular antibody recognition and orthosteric ligand binding are structurally compatible and can be accommodated together within the receptor complex.

Despite differences in ligand occupancy, all CB1 structures adopt an inactive-like transmembrane conformation consistent with previously reported inactive CB1 structures (Extended Fig. 5d-e). However, binding of Nimacimab markedly stabilizes the extracellular region of the receptor, and structural comparison further revealed that antibody engagement induces an outward displacement of the extracellular portion of TM1, accompanied by a downward shift of the membrane-proximal region of ECL2 toward the transmembrane core (Extended Fig. 5d**–**e).

Strikingly, CB1 exhibited a strong tendency to form dimeric assemblies across multiple datasets. Among them, the Nimacimab–Rimonabant–CB1 complex yielded a well-resolved dimeric structure (Fig. 2D, Extended Fig. 6a–c). The overall conformation of each protomer closely resembles that observed in the monomeric complexes, and the orthosteric ligand-binding pockets remain fully preserved (Extended Fig. 6d–e). In the dimer structure, two CB1 protomers contact mainly through TM1-TM2, while each protomer is independently engaged by one Fab fragment (Extended Fig. 6a–c). This TM1–TM2-mediated arrangement is consistent with the diverse transmembrane dimerization modes reported in class A GPCRs, in which the dimer interfaces can involve different helical surfaces, including TM1–TM2–H8^35^, TM5–TM6^36,37^ or ECL1– TM3^38^. Although the physiological relevance of this dimeric arrangement remains unclear, the observation of CB1 dimers across datasets suggests an intrinsic oligomerization propensity and provides a structural framework for understanding potential higher-order organization of CB1 receptors signaling^39,40^.

### Structural basis of Nimacimab recognition by CB1

In the cryo-EM structure of the Nimacimab–CB1 complex, the Fab fragment binds to the extracellular surface of CB1, primarily engaging ECL2 and the adjacent extracellular region of TM3, while leaving the orthosteric binding pocket accessible from the extracellular side (Fig. 3a). The antibody heavy chain mediates most of the receptor recognition, with CDRH1 and CDRH3 extensively contacting ECL2, whereas the light chain contributes limited interactions mainly through CDRL3 with residues located in ECL2 and the extracellular tip of TM3 (Fig. 3a).

**Fig. 3.**
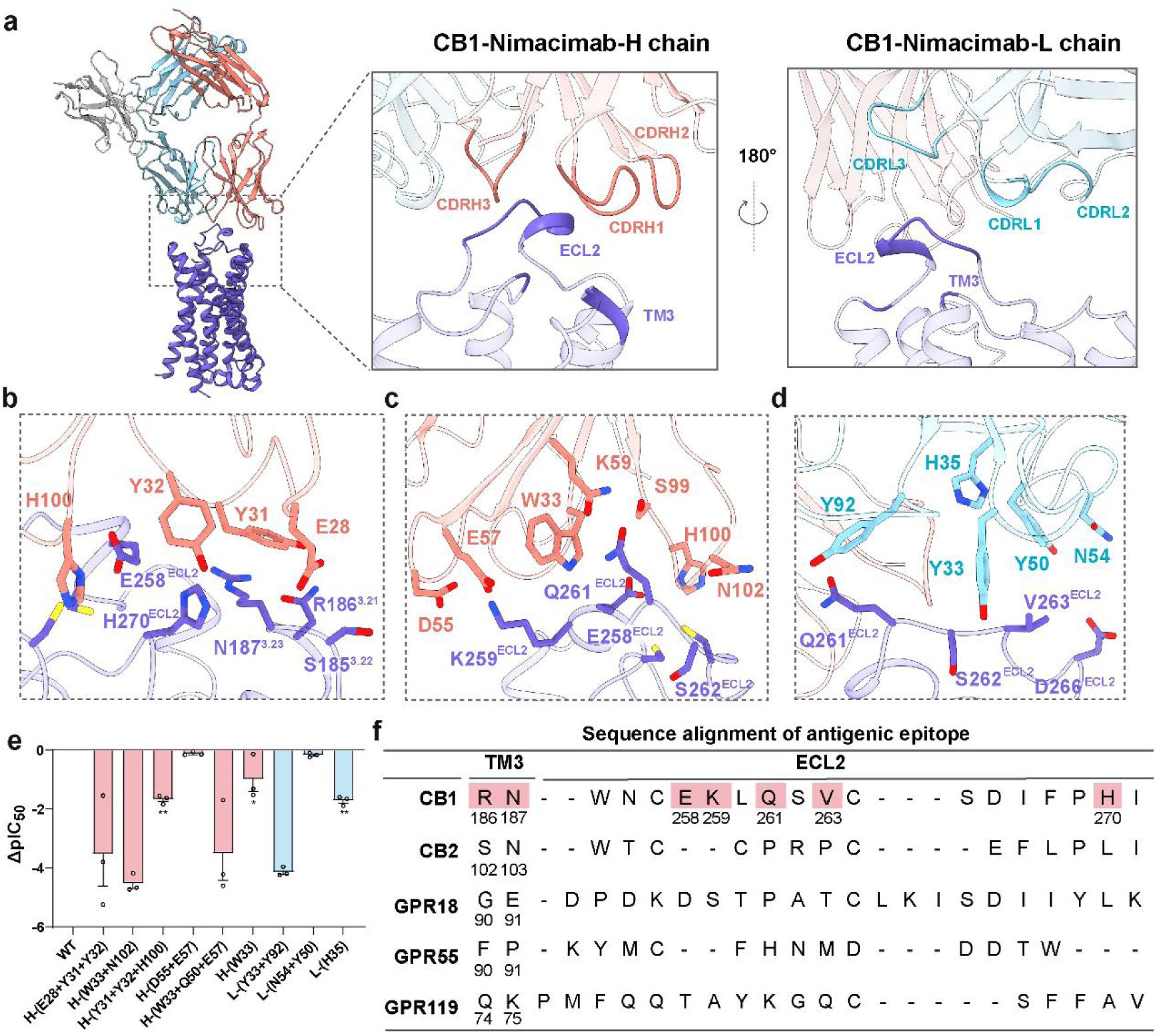
Structural basis of CB1 recognition by Nimacimab a, Overall structure and close-up views of the Nimacimab–CB1 interface. **b–d,** Detailed interactions between CB1 and the Nimacimab heavy chain**(b, c)** or light chain**(d)**. **e,** Effects of structure-guided Nimacimab mutations on CB1 inhibition, shown as ΔpIC_50_ values relative to WT Nimacimab. Negative values indicate reduced inhibitory potency. Data are mean ± s.e.m. from three independent experiments (n = 3). Statistical significance was determined using pIC_50_ values by one-way ANOVA followed by Dunnett’s multiple-comparisons test versus WT Nimacimab. *P < 0.05, **P < 0.01. Detailed values are provided in Supplementary Table 3. **f,** Sequence alignment of the Nimacimab-binding epitope across CB1, CB2, GPR18, GPR55 and GPR119. CB1 residues forming the Nimacimab-binding epitope are highlighted. CB1, purple; Fab heavy chain, salmon; Fab light chain, cyan; nanobody, gray.

An extensive polar interaction network formed between the CDRs of Nimacimab and a clustered epitope on CB1 dominates the antibody–receptor interface. Specifically, heavy-chain residues E28^CDRH1^, Y31^CDRH1^, and Y32^CDRH1^ form a polar interaction network with R186^3.22^, N187^3.23^, and H270^ECL2^ (Fig. 3b). Additional interactions are concentrated within ECL2, where Q261^ECL2^ engages W33^CDRH1^, N102^CDRH3^, and K59^CDRH2^ through a combination of polar and hydrophobic contacts, while K259^ECL2^ forms hydrogen bonds with D55^CDRH2^ and E57^CDRH2^ (Fig. 3c). Notably, W33^CDRH1^ is positioned between K259^ECL2^ and Q261^ECL2^, generating a sandwich-like packing arrangement that further stabilizes the interaction interface. H100^CDRH3^ is positioned close to E258^ECL2^ and H270^ECL2^ and further reinforces the heavy-chain interface (Fig. 3c). In addition, the light chain reinforces the antibody–receptor complex through interactions involving Y92^CDRL3^ and Y33^CDRL1^ with residues in ECL2 (Fig. 3d), thereby stabilizing both the heavy-chain conformation and the overall antibody-binding interface. Collectively, these interactions stabilize the extracellular architecture of CB1 and constrain the receptor in an inactive signaling state.

To determine whether these contacts are functionally required for Nimacimab mediated inhibition, we introduced mutations designed to disrupt key antibody–receptor interactions and assessed their effects on CB1 signaling inhibition. Substitution of interface residues within Nimacimab CDRs, which disrupt interactions with E258, K259, Q261, H270 in ECL2 and R186, N187 in TM3 of CB1, markedly reduced the inhibitory potency (Fig. 3e). In contrast, mutations at positions distal to the interface, including D55A/E57A in the heavy chain and N54A/Y50A in the light chain, produced minimal effects (Fig. 3e), indicating that the observed functional changes specifically result from disruption of the antibody–receptor interaction network.

A major advantage of therapeutic antibodies lies in their high target specificity, which is typically dictated by unique epitope composition. Consistent with this principle, Nimacimab exhibits strong selectivity toward CB1 over related cannabinoid receptors. Sequence comparison of the structural epitope revealed that several key residues involved in Nimacimab recognition, including E258^ECL2^, K259^ECL2^, Q261^ECL2^, V263^ECL2^, and H270^ECL2^, are poorly conserved in CB2 and other cannabinoid-related receptors (Fig. 3f). Together, these results define the molecular epitope of Nimacimab on CB1, and demonstrate that extracellular loop engagement is essential for antibody-mediated receptor inhibition and subtype selectivity.

### Orthosteric ligand binding and structural basis for peripherally restricted inhibition of CB1

JD5037 is a peripherally restricted analog of Rimonabant generated through modification of the C3 substituent on the pyrazole scaffold. Compared with Rimonabant, which contains a lipophilic piperidinyl carboxamide group, JD5037 retains the diaryl-substituted pyrazole core but incorporates a polar substituent composed of a valinamide moiety and a para-chlorobenzenesulfonamide moiety (Fig. 4a). These amide and sulfonamide functionalities provide additional hydrogen-bonding capacity and increase the polar surface area of JD5037, features associated with reduced CNS exposure.

**Fig. 4.**
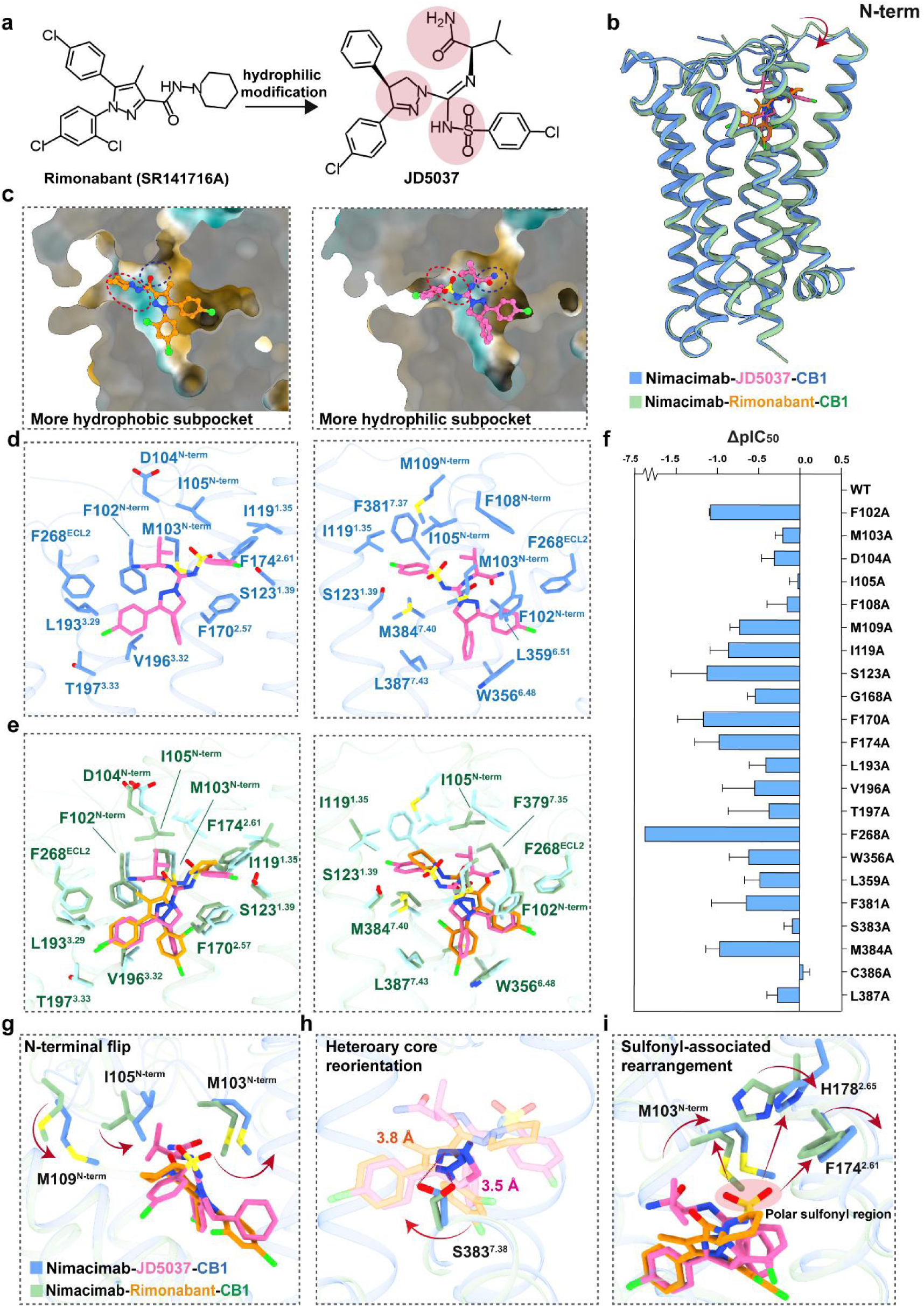
Orthosteric binding mode of JD5037 and structural basis related to peripheral restriction. a, Hydrophilic modification underlying the peripheral restriction of JD5037. **b,** Structural superposition of Nimacimab–JD5037–CB1 and Nimacimab–Rimonabant–CB1 complexes. **c,** Comparison of the extracellular ligand-binding pockets occupied by Rimonabant and JD5037. **d and e,** Detailed views of ligand recognition within the CB1 orthosteric pocket**(d).** The JD5037-bound pocket**(e)**. An overlay comparison of JD5037- and Rimonabant-bound pocket. **f,** Effects of orthosteric-pocket mutations on JD5037-mediated CB1 inhibition, shown as ΔpIC_50_ values relative to wild-type (WT) CB1. ΔpIC_50_ = pIC_50_(mutant CB1) − pIC_50_(WT CB1); negative values indicate reduced inhibitory potency. Data are mean ± s.e.m. from three independent experiments (n = 3), each performed with technical replicates. Statistical significance was determined using pIC_50_ values by one-way ANOVA followed by Dunnett’s multiple-comparisons test versus WT CB1. *P < 0.05, **P < 0.01. N.A. indicates that a reliable concentration-response fit could not be obtained. Detailed pIC_50_, Emax and P values are provided in Supplementary Table 4. **g,** Structural comparison showing the N-terminal flip associated with JD5037 binding. **h,** Reorientation of the central heteroaromatic core of JD5037 relative to Rimonabant. **i,** Sulfonyl-associated rearrangement of the extracellular pocket. JD5037, pink; Rimonabant, orange; JD5037-bound CB1, blue; Rimonabant-bound CB1, green.

To understand how this peripherally restricted antagonist engages CB1 and differs from centrally acting antagonists, we compared the cryo-EM structures of CB1 bound to JD5037 and Rimonabant. Overall, both ligands occupy the canonical orthosteric pocket formed by TM1–3 and TM6–7, with additional contributions from the N-terminus and ECL2 (Fig. 4b). Despite this shared binding mode, structural comparison revealed notable differences in ligand conformation and extracellular pocket organization. In particular, JD5037 induces a pronounced rearrangement of the extracellular vestibule, including a conformational flip of the N-terminal loop and the formation of a more hydrophilic ligand-binding environment relative to the Rimonabant-bound state (Fig. 4c).

Detailed analysis of the JD5037-bound structure revealed that, the phenyl group inserts into a hydrophobic cavity lined by F170^2.57^, M384^7.40^, and L387^7.43^, forming extensive hydrophobic and aromatic interactions analogous to those observed for Rimonabant. Meanwhile, the para-chlorophenyl group occupies a second hydrophobic sub-pocket formed by F268^ECL2^, L193^3.29^, V196^3.32^, and W356^6.48^, where it is stabilized primarily through van der Waals and aromatic-hydrophobic interactions (Fig. 4d). In addition to these two hydrophobic substituents, the valinamide and para-chlorobenzenesulfonamide groups provide additional contacts at the extracellular side of the binding pocket. The valinamide group extends toward the receptor N terminus and is positioned near F102^N-term^, M103^N-term^, D104^N-term^, F108^N-term^, and F268^ECL2^. This region forms a continuous contact interface with the surrounding extracellular residues, in which the valine side chain mainly contributes hydrophobic contacts, whereas the amide moiety makes close van der Waals contacts with F102^N-term^. In contrast, the para-chlorobenzenesulfonamide group occupies a more polar extracellular extension of the orthosteric pocket. Its para-chlorophenyl ring is embedded in an aromatic hydrophobic environment formed by F174^2.61^ and F381^7.37^, while the sulfonamide oxygen and nitrogen atoms are positioned to form hydrogen-bonding interactions with S123^1.39^ or nearby backbone atoms. Thus, this substituent combines aromatic hydrophobic anchoring with local polar coordination, making it a key recognition element for JD5037 binding. Consistent with these structural observations, mutagenesis analysis demonstrated that residues shaping both the hydrophobic core and the extracellular polar vestibule contribute to JD5037 recognition and inhibition (Fig. 4f).

Compared with Rimonabant pocket, the valinamide and sulfonamide-containing substituents of JD5037 extend toward the extracellular entrance of the binding pocket and engage residues from the receptor N-terminus and ECL2, thereby creating a localized polar environment in the extracellular region of the orthosteric cavity. This redistribution of polarity induces substantial rearrangements of surrounding hydrophobic residues, including M103^N-term^, I105^N-term^, and M109^N-term^, ultimately driving a conformational flip of the N-terminal loop and reshaping the extracellular vestibule surrounding the ligand (Fig. 4g). Additional structural rearrangements are observed near the extracellular end of TM2, where the sulfonamide-containing substituent induces conformational changes in F174^2.61^ and H178^2.65^ to accommodate the more polar extracellular extension of JD5037 (Fig. 4i). Furthermore, comparison of the central heteroaromatic scaffold revealed altered interactions with S383^7.38^, accompanied by local reorientation of the ligand core relative to Rimonabant (Fig. 4h). This altered interaction pattern distinguishes JD5037 from centrally acting antagonists such as Rimonabant and provides a structural basis for understanding how targeted chemical modifications of a shared scaffold can influence peripheral versus central pharmacological properties.

### Cooperative inhibition of CB1 by extracellular antibody engagement and orthosteric antagonism

To further understand the inhibitory mechanism of Nimacimab, we first investigated how extracellular antibody binding influences CB1 conformational dynamics. In the Nimacimab-bound structures, the antibody engages ECL2 and adjacent extracellular regions while leaving the orthosteric pocket accessible, suggesting that receptor inhibition is achieved through allosteric stabilization of the extracellular architecture rather than direct competition with orthosteric ligands. Consistent with this model, molecular dynamics simulations revealed markedly reduced conformational fluctuations in ECL2 upon antibody binding (Fig. 5a). Stabilization of the extracellular loop region promoted tighter packing of F268^ECL2^ and neighboring hydrophobic residues, leading to formation of a more compact aromatic interaction core (Fig. 5b-c). This conformational compaction was reflected by a concentrated radius of gyration (Rg) distribution centered near 6.5 Å, whereas the receptor-only system displayed an additional population at Rg > 7 Å, indicative of a more dynamic and expanded extracellular core (Extended Fig. 7a).

**Fig. 5.**
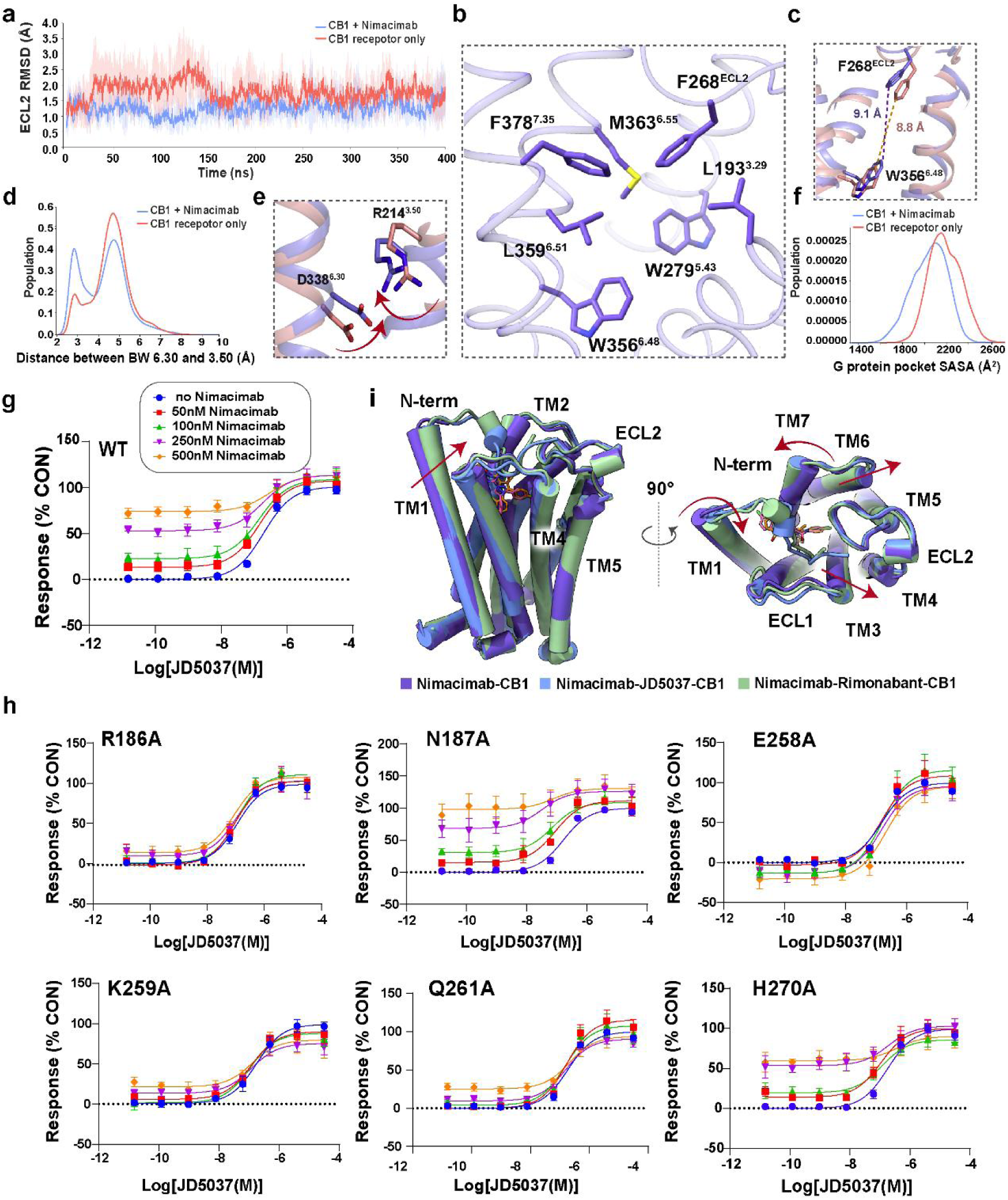
Mechanism of Nimacimab-mediated CB1 inhibition and its cooperative effect with JD5037. a, Time course of ECL2 RMSD in the CB1 receptor-only and CB1–Nimacimab systems. Solid lines represent mean values, and shaded regions indicate s.d. **b,** Close-up view of the hydrophobic pocket formed by ECL2 in a representative CB1-Nimacimab structure. Key residues are shown as sticks. **c,** Comparison of the minimal distance between F268^ECL2^ and W356^6.48^ in representative structures from the CB1 receptor-only and CB1-Nimacimab systems. **d,** Distribution of the distance between D338^6.30^ and R214^3.50^ in the CB1 receptor-only and CB1-Nimacimab systems. **e,** Close-up view illustrating the relative movements of D338^6.30^ and R214^3.50^ in the CB1 receptor-only and CB1-Nimacimab systems. Arrows indicate the direction of motion. **f,** Distribution of the solvent-accessible surface area of the G protein-binding pocket in the CB1 receptor-only and CB1–Nimacimab systems. **g,** JD5037 concentration-response curves in wild-type CB1 in the absence or presence of increasing concentrations of Nimacimab. **h,** JD5037 concentration-response curves for CB1 epitope mutants in the absence or presence of Nimacimab. Data are mean ± sem from three independent experiments **i,** Structural superposition of Nimacimab–CB1, Nimacimab–JD5037–CB1, and Nimacimab–Rimonabant–CB1 complexes, shown from side and extracellular views. For MD analyses, receptor-only CB1 is shown in red and CB1–Nimacimab in blue. For structural comparisons, Nimacimab–CB1 is shown in purple, Nimacimab–JD5037–CB1 in blue and Nimacimab–Rimonabant–CB1 in green. Functional data in g,h are mean ± s.e.m. from three independent experiments (n = 3), each performed with technical replicates.

These extracellular rearrangements propagated into the transmembrane region and stabilized canonical inactive-state micro-switches of CB1. In particular, antibody binding shifted the PIF motif toward an inactive-like geometry, as evidenced by increased values of the composite distance metric d(3.40–6.48) + d(5.51–6.44), with the receptor-only and CB1–Nimacimab simulations shown in red and blue, respectively(44). (Extended Fig. 7b-c). Concomitantly, the intracellular ionic lock region became further stabilized, with the antibody-bound ensemble preferentially populating a short TM3-TM6 distance near ∼3 Å (Fig. 5d). A representative structural view further illustrates the relative positioning of D338^6.30^ and R214^3.50^ in the intracellular ionic-lock region (Fig. 5e). Propagation of these allosteric constraints ultimately resulted in contraction of the intracellular G protein-binding cavity, reflected by decreased solvent-accessible surface area (SASA) of intracellular pocket residues in the antibody-bound state (Fig. 5f). Together, these findings support a model in which Nimacimab functions as an extracellular inverse agonist that suppresses CB1 signaling by constraining extracellular conformational dynamics and allosterically stabilizing the receptor in a low-activity state.

Given that Nimacimab and JD5037 bind to spatially distinct regions of CB1, we next asked whether extracellular antibody engagement could functionally cooperate with orthosteric antagonism. To address this question, JD5037 concentration-response curves were measured in the absence or presence of increasing concentrations of Nimacimab. In wild-type CB1, Nimacimab produced a concentration-dependent upward shift of the lower plateau of the JD5037 response curve, with the fitted bottom increasing from ∼0.24% in the absence of antibody to 17.67%, 27.52%, 57.74%, and 83.02% in the presence of 50, 100, 250, and 500 nM Nimacimab, respectively (Fig. 5g). By comparison, the maximal response and apparent potency of JD5037 changed only modestly. Consequently, the response span was progressively reduced, indicating that Nimacimab partially suppresses CB1 signaling on its own while remaining fully compatible with additional orthosteric inhibition by JD5037.

To determine whether this cooperative effect depends on the structurally defined antibody epitope, alanine substitutions were introduced at key CB1 residues involved in Nimacimab recognition. These mutations did not substantially alter the concentration-response profile of JD5037 alone, indicating that the orthosteric antagonist-binding pocket remained functionally intact. However, mutations at residues within the antibody-binding interface, particularly R186^3.22^, E258^ECL2^, K259^ECL2^, and Q261^ECL2^, markedly attenuated the Nimacimab-dependent elevation of the lower plateau (Fig. 5h). These findings directly link the enhanced inhibitory response to antibody engagement of the extracellular TM3–ECL2 region.

Superposition of the three Nimacimab-bound CB1 structures showed that antibody engagement is fully compatible with distinct orthosteric antagonists, while the presence of small molecule ligands further stabilizes a more compact inactive-like receptor architecture (Fig. 5i). Consistent with this observation, MD simulations revealed that the ligand-bound receptor exhibited a narrower ECL2 RMSD distribution compared with the antibody-only system, indicating that orthosteric ligand binding further restricts extracellular loop conformational flexibility (Extended Fig. 7d). Structural measurements of the intracellular TM5–TM6 cavity across the Nimacimab-only and ternary complexes further demonstrated that addition of the orthosteric antagonist promotes additional contraction of the intracellular G protein-binding pocket (Extended Fig. 7e-g). Together, these findings support a model in which Nimacimab and JD5037 inhibit CB1 through complementary extracellular and orthosteric mechanisms.

## Discussion

The cannabinoid receptor CB1 remains an important therapeutic target for obesity and metabolic disorders, yet the clinical failure of centrally acting antagonists such as Rimonabant highlighted the challenge of separating metabolic efficacy from CNS-associated adverse effects^10–12^. Here, through structural, pharmacological, mutational, and molecular dynamics analyses, we define two mechanistically distinct yet functionally cooperative modes of peripheral CB1 inhibition mediated by extracellular antibody engagement and orthosteric antagonism.

A central finding of this study is the identification of Nimacimab as an extracellular inverse agonist of CB1. Unlike classical cannabinoid antagonists that directly occupy the orthosteric pocket, Nimacimab binds to ECL2 and adjacent extracellular regions without occluding the ligand-binding cavity. Antibody engagement stabilizes a distinct inactive-like extracellular architecture, restricts extracellular conformational dynamics, and allosterically propagates inactive-state constraints into the transmembrane core. MD simulations further revealed stabilization of inactive-state microswitches and reduced accessibility of the intracellular G protein-binding cavity. Together, these findings establish a previously uncharacterized mechanism of GPCR inhibition that can be conceptually defined as extracellular allosteric inverse agonism.

Importantly, this inhibitory mechanism differs fundamentally from previously characterized GPCR-targeting antibodies, many of which function through steric blockade of ligand access^24^ or direct occupation of extracellular portions of the orthosteric site^24,41,42^. In contrast, Nimacimab remains fully compatible with simultaneous orthosteric ligand binding and suppresses receptor signaling through extracellular conformational stabilization and long-range allosteric regulation. These findings expand the conceptual framework for GPCR inhibition by demonstrating that extracellular regions can function as active conformational control elements capable of governing receptor signaling states.

Our structures also provide insight into the mechanism underlying peripheral restriction of JD5037. Although JD5037 preserves the conserved orthosteric binding mode of classical CB1 inverse agonists, incorporation of valinamide and sulfonamide-containing substituents redistributes interaction density toward extracellular and more solvent-accessible regions of the receptor. This shift generates a localized polar environment within the extracellular vestibule and induces coordinated rearrangements of the N-terminus, ECL2, and TM2, suggesting a structural basis by which increased extracellular polarity may contribute to peripheral restriction while maintaining receptor affinity. Notably, this extracellularly extended binding mode remains fully compatible with Nimacimab engagement, enabling cooperative inhibition of CB1 through simultaneous stabilization of both the extracellular vestibule and the orthosteric transmembrane core. Together, these findings highlight the importance of long-range allosteric communication between extracellular loops and the transmembrane core in regulating CB1 activation dynamics.

In summary, our study establishes the structural basis of extracellular allosteric inverse agonism by Nimacimab, elucidates the molecular architecture of peripheral orthosteric antagonism by JD5037, and reveals a cooperative inhibitory mechanism between extracellular antibody engagement and orthosteric ligand binding. Together, these findings provide a structural framework for the development of next-generation peripherally selective CB1 therapeutics.

## Acknowledgements

This work was supported by grants from the National Natural Science Foundation of China (32501066 to H.L., 32130022, 82495184, 82121005 to H.E.X.; W2412030 to F.X.); the National Key R&D Program of China (2022YFA1302900 to W.Y., 2022YFC2703105 to H.E.X.); the National Key R&D Program “Strategic Scientific and Technological Innovation Cooperation” Key Project (2022YFE0203600), released by the Ministry of Science and Technology; the Shanghai Municipal Science and Technology Major Project (2019SHZDZX02 to H.E.X.); the Shanghai Municipal Science and Technology Major Project (H.E.X.); and the Strategic Priority Research Program of the Chinese Academy of Sciences (XDB0830000, XDA0530000, XDB37030103 to H.E.X.); Shanghai Key Laboratory for Antibody-Drug Conjugates with Innovative Target (24dz2261300); the Natural Science Foundation of Shanghai (25ZR1402552 to H.L.). F.X. is a SANS Exploration Scholar. The cryo-EM data were collected at the Shanghai Advanced Electron Microscope Center, Shanghai Institute of Materia Medica and Shanghai Tech University. We thank Q. Yuan, K. Wu, W. Hu, C. Liu, S. Li, and S. Zhang for providing technical support and assistance during data collection at the Shanghai Advanced Electron Microscope Center, Shanghai Institute of Materia Medica.

## Author contributions

H.E.X., F.X. and H.L. initiated the project. Y.G. and J.W. screened the expression constructs and optimized the purification conditions of protein complexes, prepared protein samples for cryo-EM grid preparation, and participated in figure preparation. Y.L. performed the cell-based functional assays and participated in figure preparation. Y.B., X.Z. and Y.T. participated in functional assays. H.L. and Y.Y. performed the cryo-EM data calculations, model building. H.S. and J.H. performed the MD simulations and participated in figure preparation. X.H. and J.L. assisted in the methodology discussion of the MD simulations. W.H. screened the EM sample and collected EM movies. H.L. wrote the draft and Y.G. prepared the figures. H.E.X. and F.X. revised the manuscript. H.E.X., F.X. and H.L. conceived and supervised the project and wrote the manuscript with inputs from the authors.

## Data, Materials, and Software Availability

The cryo-EM density maps and corresponding atomic coordinates reported in this study have been deposited in the Electron Microscopy Data Bank and the Protein Data Bank under accession codes. EMD-80863 and PDB 26TK for the Nimacimab–CB1 complex, EMD-80862 and PDB 26TJ for the Nimacimab–JD5037–CB1 complex, EMD-80555 and PDB 26DG for the monomeric CB1–Nimacimab–Rimonabant complex, and EMD-80556 and PDB 26DH for the dimeric CB1–Nimacimab–Rimonabant complex. All other data supporting the findings of this study are available within the article and its Supplementary Information or from the corresponding authors upon reasonable request. Plasmids and additional materials generated in this study will be made available from the corresponding authors upon reasonable request. Cryo-EM data processing was performed using cryoSPARC. Model building was carried out in Coot, UCSF ChimeraX, and ISOLDE. Real-space refinement was performed in PHENIX. Graphs and statistical analyses were generated using GraphPad Prism.

## Methods

### Constructs

For the Nimacimab–CB1 and Nimacimab–JD5037–CB1 complexes, the human CB1 construct was cloned into a pcDNA3.1-derived mammalian expression vector. The construct contained CB1 residues 77–306 and 335–402, with thermostabilized BRIL inserted into the third intracellular loop (ICL3). BRIL was connected to TM5 and TM6 through short engineered linkers, RQL and ERA, at the TM5–BRIL and BRIL–TM6 junctions, respectively. Four thermostabilizing mutations, T210A, E273K, T283V, and R340E, were introduced to improve receptor stability and expression^43^. A TGP tag followed by tandem AviTag–Strep-tag II–FLAG tags was fused to the C terminus for purification.

For the Nimacimab–Rimonabant–CB1 complex, the gene encoding human CB1 was cloned into the pFastBac1 vector. The construct contained CB1 residues 71–304 and 333–425, with thermostabilized BRIL inserted into ICL3. The thermostabilizing mutations T210A, E273K, T283V, and R340E were introduced to improve receptor stability^43^. BRIL was flanked by ARRQL and ERARSTL linkers at the TM5–BRIL and BRIL–TM6 junctions^44^, respectively. A C-terminal 10×His tag was used for purification.

The heavy and light chains of Nimacimab Fab were cloned into pcDNA3.0-derived mammalian expression vectors and pFastBac Dual insect cell expression vectors. For mammalian expression, the heavy chain was truncated at residue 223, whereas the light chain was fused to a C-terminal 8×His tag. For insect cell expression, the heavy chain was truncated at residue 215 and fused to a C-terminal 6×His tag. The anti-BRIL Fab was cloned into the pFastBac Dual vector with a C-terminal 6×His tag.

## Cell Expression

### Mammalian Cell Expression

For mammalian expression of CB1 and Nimacimab Fab, suspension HEK293E cells (Life Technologies) were cultured in SMM 293T-II medium (Sino Biological Inc. #A14635) at 37 °C under 5% CO_2_ shaking at 130 rpm in a Multitron-Pro shaker. Cells were grown to a density of approximately 2.5–3.0 × 10^6^ cells/mL and transiently transfected using polyethyleneimine (PEI; Yeasen, #40816ES03). Sodium butyrate (Sinopharm Chemical Reagent Co., Ltd.) was added to a final concentration of 10 mM at 8–12 h after transfection to enhance protein expression. After 48–72 h, cultures expressing CB1 were harvested by centrifugation, and the resulting cell pellets were stored at −80 °C until purification. For Nimacimab Fab expression, clarified culture supernatants were collected for subsequent purification.

### Insect Cell Expression

For insect cell expression, the modified CB1 construct and Nimacimab Fab were expressed in Trichoplusia ni Hi5 insect cells using the Bac-to-Bac baculovirus expression system (Invitrogen, #B85502). Cells were infected with recombinant baculoviruses at a density of 2 × 10^6^ cells/mL and cultured at 27 °C for 48 h after infection. For CB1 expression, cells were collected by centrifugation, and the resulting cell pellets were stored at −80 °C until purification. For secreted Nimacimab Fab expression, culture supernatants were collected by centrifugation and stored at 4 °C before purification.

### Purification of Nimacimab Fab

Culture supernatants containing secreted Nimacimab Fab were collected by centrifugation at 2000 rpm for 20 min at room temperature to remove cells and large debris. The clarified supernatants were adjusted to pH 8.0 with 1 M Tris and sequentially supplemented with NiCl_2_, CaCl_2_, and MgCl_2_ to final concentrations of 1 mM, 5 mM, and 10 mM, respectively. The mixture was then incubated with gentle stirring at room temperature for 1.5 h, followed by centrifugation at 8,000 rpm for 30 min at 4 °C to remove precipitates. The resulting supernatant was loaded by gravity flow onto a column packed with 2.5 mL nickel-affinity resin (NEBExpress®, NEB #S1427). The column was washed with 20 column volumes of wash buffer containing 20 mM HEPES, pH 7.4, 300 mM NaCl, 25 mM imidazole, and 10% (v/v) glycerol, and bound Nimacimab Fab was eluted with 20 mM HEPES, pH 7.4, 300 mM NaCl, 250 mM imidazole, and 10% (v/v) glycerol.

Eluted fractions containing Nimacimab Fab were pooled and concentrated using a 30-kDa MWCO centrifugal concentrator at 4 °C. The concentrated protein was filtered through a 0.22-μm membrane and further purified by size-exclusion chromatography on a Superdex 75 10/300 GL column equilibrated with 20 mM HEPES, pH 7.4, and 125 mM NaCl. Fractions corresponding to monomeric Nimacimab Fab were pooled and concentrated to approximately 33 mg/mL. Protein purity was assessed by SDS-PAGE. The purified Fab was supplemented with 10% glycerol, flash-frozen in liquid nitrogen, and stored at −80 °C until use.

### Purification of Nimacimab–CB1 and Nimacimab–JD5037–CB1 complexes

Cell pellets expressing CB1 were resuspended in ice-cold buffer containing 20 mM HEPES, pH 7.4, 300 mM NaCl, protease inhibitor cocktail (SparkJade, China, #SJ-MK0001-D), and 0.1 mM TCEP (Sigma-Aldrich, #646547-10X1ML). Cell membranes were isolated by ultracentrifugation at 30,000 rpm for 45 min at 4 °C. The membrane pellets were resuspended in the same buffer and solubilized with 0.5% (w/v) DDM (Anatrace, #D310) and 0.1% (w/v) CHS at 4 °C for 3 h. Insoluble material was removed by ultracentrifugation at 30,000 rpm for 45 min at 4 °C, and the supernatant was incubated with anti-FLAG resin (GenScript, #L00432-25) for 2 h at 4 °C.

The resin was washed with buffer containing 20 mM HEPES, pH 7.4, 150 mM NaCl, 0.06% (w/v) LMNG (Anatrace, #NG310) and 0.02% (w/v) GDN (Anatrace, #GDN101). Bound CB1 was eluted with the same buffer supplemented with 200 ng/μL FLAG peptide (GenScript, #RP10586-1).

For assembly of the Nimacimab–CB1 complex, purified CB1 was concentrated and incubated with Nimacimab Fab at a molar ratio of CB1:Nimacimab Fab = 1:1.2 on ice for at least 30 min. For assembly of the Nimacimab–JD5037–CB1 complex, purified CB1 was incubated with 10 μM JD5037, Nimacimab Fab, and anti-Fab nanobody at a molar ratio of CB1: Nimacimab Fab: anti-Fab nanobody = 1:1.2:1.5 on ice for 30 min.

The assembled complexes were further purified by size-exclusion chromatography on a Superdex 200 Increase 10/300 GL column (GE Healthcare). The CB1–Nimacimab complex was purified in buffer containing 20 mM HEPES, pH 7.4, 150 mM NaCl, 0.00075% (w/v) LMNG, 0.00025% (w/v) GDN and 0.0002% (w/v) CHS. The Nimacimab–JD5037–CB1 complex was purified in buffer containing 20 mM HEPES, pH 7.4, 125 mM NaCl, 10 μM JD5037 (TargetMol, #T4453), 0.00075% (w/v) LMNG, 0.00025% (w/v) GDN and 0.0002% (w/v) CHS. Fractions corresponding to the monodisperse complex peak were pooled and concentrated for subsequent cryo-EM grid preparation.

### Purification of Nimacimab–Rimonabant–CB1 complex

The cell pellet expressing CB1 was thawed on ice and lysed in a hypotonic buffer containing 10 mM HEPES pH 7.5, 10 mM MgCl_2_, 20 mM KCl, and a protease inhibitor cocktail (MedChemExpress, #HY-K0010) by dounce homogenization. 10 µM Rimonabant was added into the suspension. After centrifugation at 140,000 × g for 40 min at 4 °C, the resulting pellet was resuspended and lysed in a hypertonic buffer containing 10 mM HEPES pH 7.5, 10 mM MgCl_2_, 20 mM KCl, 1 M NaCl, and a protease inhibitor cocktail (MedChemExpress, #HY-K0010) by dounce homogenization. Rimonabant was added to a final concentration of 10 µM, and the suspension was centrifuged at 140,000 × g for 40 min at 4 °C. The solubilized supernatant was collected and incubated with Talon super-flow metal affinity resin (Clontech, #635670) with 20 mM imidazole added, followed by an overnight incubation at 4 °C. The mixture was then loaded onto a chromatography column and washed with 14 column volumes (CV) of washing buffer containing 25 mM HEPES pH 7.5, 100 mM NaCl, 10% glycerol, 0.1% (w/v) LMNG, 0.02% (w/v) CHS, 30 mM imidazole, and 10 µM Rimonabant. The mixture was then washed with 14 column volumes (CV) of washing buffer containing 25 mM HEPES pH 7.5, 100 mM NaCl, 10% glycerol, 0.1% (w/v) LMNG, 0.03% (w/v) CHS, 50 mM imidazole, and 10 µM Rimonabant. Elution of the complex was performed with 6 CV of elution buffer containing 25 mM HEPES pH 7.5, 100 mM NaCl, 10% glycerol, 0.01% (w/v) LMNG, 0.002% (w/v) CHS, 250 mM imidazole, and 50 µM Rimonabant. For assembly of the CB1–Nimacimab–Rimonabant complex, purified CB1–Rimonabant was concentrated to 0.5 mL and incubated with Nimacimab Fab, anti-BRIL Fab, and anti-Fab nanobody at a molar ratio of CB1:Nimacimab Fab:anti-BRIL Fab:anti-Fab nanobody = 1:2:1.25:1.25, followed by an overnight incubation at 4 °C. The assembled complexes underwent size-exclusion chromatography on a Superdex 200 10/300 GL column (GE Healthcare, #28990944) with running buffer containing 20 mM HEPES pH 7.5, 100 mM NaCl, 0.00075% (w/v) LMNG, 0.0002% (w/v) CHS, and 2 µM Rimonabant. The peak fractions were analyzed by SDS-PAGE and concentrated to a final protein concentration for electron microscopy experiments.

### Cloning, expression and purification of anti-BRIL Fab

The anti-BRIL Fab was cloned into the pFastBac Dual vector with a C-terminal 6×His tag and expressed in secreted form using the baculovirus expression system in Trichoplusia ni Hi5 insect cells^45^. Cells were grown to a density of 2 × 10^6^ cells/mL and infected with recombinant baculovirus at an MOI of 5. Culture supernatants were collected by centrifugation at 2,000 × g for 30 min at 48 h after infection.

The clarified supernatant from 1 L of culture was loaded onto 2 mL Ni-NTA resin. The resin was washed with 15 column volumes of wash buffer containing 20 mM Tris-HCl, pH 7.55, 150 mM NaCl, and 20 mM imidazole. Bound protein was eluted with the same buffer supplemented with 250 mM imidazole. Eluted fractions were further purified by size-exclusion chromatography on a Superdex 200 10/300 GL column equilibrated with 20 mM Tris-HCl, pH 7.55, 100 mM NaCl, and 10% glycerol. Fractions containing anti-BRIL Fab were pooled, concentrated to 4 mg/mL using a 50-kDa MWCO concentrator, flash-frozen in liquid nitrogen, and stored at −80 °C until use.

### Cryo-EM Grid preparation and Data Acquisition

Three microliters of the purified Nimacimab–CB1, Nimacimab–JD5037–CB1, and Nimacimab–Rimonabant–CB1 complexes at concentrations of 6.8 mg/mL, 9.2 mg/mL, and 4.3 mg/mL, respectively, were applied to glow-discharged Quantifoil Au 300-mesh gold grids. The samples were blotted for 3–5 s under 100% humidity at 4 °C and vitrified by plunging into liquid ethane using a Vitrobot Mark IV (Thermo Fisher Scientific).

Cryo-EM data were collected on a 300-kV FEI Titan Krios electron microscope. Movies for the CB1–Nimacimab and CB1–Nimacimab–JD5037 complexes were recorded using a Falcon 4i direct electron detector. Movies for the CB1–Nimacimab– Rimonabant complex were recorded using a Gatan K3 direct electron detector. For the Nimacimab–CB1 and Nimacimab–JD5037–CB1 complexes, movies were recorded at a nominal magnification of 105,000×, corresponding to a calibrated pixel size of 0.73 Å, with a defocus range of −1.0 to −2.0 μm and a total electron exposure of 50 e^−^/Å². For the Nimacimab–Rimonabant–CB1 complex, movies were recorded at a nominal magnification of 105,000×, corresponding to a calibrated pixel size of 0.832 Å, with a defocus range of −1.0 to −2.5 μm and a total electron exposure of 60 e^−^/Å². Detailed data collection and processing statistics are summarized in Supplementary Table 1.

### Cryo-EM Data Processing

Collected movies were processed in cryoSPARC^46^. Raw movie frames were first subjected to motion correction, followed by contrast transfer function (CTF) estimation using Patch CTF Estimation. Micrographs with poor ice quality, severe drift, or abnormal CTF parameters were discarded before particle picking. Particles were automatically picked, extracted, and subjected to multiple rounds of 2D classification to remove contaminants, ice-derived artifacts, damaged particles, and poorly aligned projections. Selected particles were used for ab initio reconstruction and subsequent 3D classification. Well-defined particle subsets were further refined by homogeneous refinement and non-uniform refinement to generate the final reconstructions. Local resolution estimation and map sharpening were performed for the final maps in cryoSPARC^47^. Overall map resolutions were determined according to the gold-standard Fourier shell correlation (FSC) criterion at 0.143.

### Model Building and Refinement

Initial atomic models were generated by docking the inactive-state CB1 crystal structure (PDB: 5TGZ)^43^ into the cryo-EM maps using UCSF ChimeraX^48^. The models were then manually adjusted in Coot^49^ according to the density features and further optimized in ISOLDE^50^ where needed. Final real-space refinement was carried out using PHENIX 1.18^51^. Model quality was evaluated using standard stereochemical criteria, including Ramachandran statistics, clash score, and bond geometry. The final refined models showed good agreement with the experimental density maps.

## Functional Assays

### GloSensor cAMP Assay

HEK293T cells were transiently cotransfected with human CB1 receptor and GloSensor-22F plasmids. After 24 h, cells were harvested, resuspended in CO_2_-independent medium at 6 × 10^5^ cells/mL, and plated in white 384-well plates at 6,000 cells per well. GloSensor reagent (Promega, #E1290) was added in an equal volume, and plates were incubated for 1 h at 37 °C in 5% CO_2_. For antagonism assays, serial dilutions of JD5037, Rimonabant, or Nimacimab were added with 2.5 μM forskolin. For combinatorial antagonism assays, cells were stimulated with AZD1940 and forskolin in the presence of Nimacimab at final concentrations of 0, 50, 100, 250, or 500 nM, followed by addition of JD5037 at the indicated concentrations. Luminescence was measured immediately after compound addition using an EnVision plate reader (PerkinElmer). Data were analyzed by nonlinear regression in GraphPad Prism.

### NanoBiT β-arrestin recruitment assay

β-Arrestin1/2 recruitment was measured using a NanoBiT assay. Human CB1 was C-terminally fused to LgBiT through a 5GSA linker, and β-arrestin1/2 was N-terminally fused to SmBiT using the same linker. The pBiT constructs were co-transfected into HEK293T cells at a 1:1 ratio. After 24 h, cells were harvested, resuspended in PBS, and seeded into white 384-well plates at 6,000 cells per well in 10 μL. Coelenterazine substrate (Yeasen, #40904ES02) was added and equilibrated before ligand addition. JD5037, Nimacimab, or Rimonabant was added in the presence of AZD1940 to measure inhibition of AZD1940-induced β-arrestin1/2 recruitment. Luminescence was recorded using an EnVision plate reader (PerkinElmer).

## Statistical analysis

All cell-based functional experiments were independently repeated at least three times unless otherwise indicated. For GloSensor cAMP assays, each concentration was tested in technical triplicates in each independent experiment. Technical triplicates were averaged within each independent experiment before statistical analysis. Raw luminescence values were normalized to the corresponding within-plate controls, as indicated in the figure legends, and concentration–response curves were fitted by nonlinear regression in GraphPad Prism. IC_50_ values were converted to pIC_50_ values using −log_10_(IC_50_), with IC_50_ expressed in molar units. Data are presented as mean ± SEM from independent experiments.

For individual inhibition assays, the inhibitory activities of JD5037, Rimonabant, and Nimacimab were evaluated based on their suppression of agonist-induced or basal CB1 signaling under the indicated assay conditions. For co-treatment experiments, the effect of Nimacimab on JD5037-mediated CB1 inhibition was assessed by comparing fitted bottom values, pIC_50_ values, and response spans across different Nimacimab concentrations. For β-arrestin recruitment assays, luminescence signals were normalized to the AZD1940-induced response and analyzed by nonlinear regression in GraphPad Prism.

Comparisons between two groups were performed using an unpaired two-tailed Student’s t test. Comparisons among multiple groups were performed using one-way ANOVA followed by Dunnett’s multiple-comparisons test versus the corresponding WT or control group, as appropriate. N.A. indicates that a reliable concentration– response fit could not be obtained by nonlinear regression in GraphPad Prism. A value of P < 0.05 was considered statistically significant.

## MD simulations

We performed MD simulations of CB1 receptor-only and CB1-Nimacimab complex. CB1 receptor-only was constructed by removing Nimacimab from our complex. Systems were embedded into a 75 Å × 75 Å POPC lipid bilayer via CHARMM-GUI server^52–54^, surrounded by a 22.5 Å aqueous layer. Ionic strength was adjusted to 0.15 mol/L NaCl plus counterions. CHARMM36m all-atom force field was employed for each system^55^. Systems underwent a minimization and a six-step equilibration process following the prep protocol^56^. Three independent 400 ns production runs were conducted for each system using pmemd.cuda in Amber22^57^ under the NPT ensemble at temperature of 303.15 K and a pressure of 1 atm. Hydrogen mass repartitioning (HMR) were used to accelerate the simulations with a 4 fs time step^58^ Long-range electrostatic interactions were managed using the Particle Mesh Ewald method, while short-range electrostatic and van der Waals interactions were handled with a 12 Å cutoff, transitioning smoothly between 10 Å and 12 Å. SHAKE was applied to constrain the bonds containing hydrogens^59^ RMSD, minimal distances, SASA, and representative structures were analyzed using CPPTRAJ^60^. The G protein binding pocket is defined as the residues whose minimal distance is shorter than 4.5 Å in CB1-G protein complex (PDB ID: 8K8J).

